# The opossum and other didelphids in the cultures of Costa Rican society and indigenous peoples: an approach to ecology and conservation

**DOI:** 10.1101/2024.01.02.573906

**Authors:** Yara Azofeifa Romero, Francisco Durán Alvarado

## Abstract

The opossum *Didelphis marsupialis*, commonly referred to as the *zorro pelón* in Costa Rica, is a marsupial that holds significant representation in both Costa Rican and indigenous cultures, such as the Talamanca natives. This species, along with the other ten opossums found in Costa Rica, constitute one of the groups of mammals with the least representation in scientific literature. This study aims to (1) document the knowledge, uses, perceptions, and human-*zorro pelón* interactions through surveys conducted with individuals in three types of zones: rural, semi-urban, and urban; and (2) explore the presence of opossums in the 19th-century worldview of farmers and in the cosmovision of indigenous peoples. We performed an analysis on the text and Likert-scale data collected from 296 surveys conducted across seven provinces in Costa Rica. Nonparametric tests were conducted to assess whether the type of zone influenced the interactions and perceptions of individuals. Valuable information was obtained from key participants and informants regarding the behavior, observation sites, diet, and population trends of the species. The remarkable adaptability of the *zorro pelón* is evident in rural and semi-urban environments, but not in urban zones. The human-*zorro pelón* interaction data between the rural and semi-urban zones are similar, but these differ significantly from the urban zone. Although people surveyed generally have a positive perception of the opossum, there has been a notable devaluation over time when compared to the imaginary of indigenous cultures. Parallels were observed in the beliefs and uses of marsupials between pre-Columbian societies and contemporary Costa Rican societies. As part of the educational strategies for the conservation of marsupials, it is important to promote their appreciation and refute negative stereotypes, such as by informing about their ecological role and their ability to resist pathogens and venom. In this context, the rich symbolism of the opossum can be utilized to adapt the message to the various regions of America.

## INTRODUCTION

The opossum, or *zorro pelón* (*Didelphis marsupialis*), is one of the ten didelphids present in Costa Rica; opossum is also known as *zarigüeya* in Spanish. (Wainwright, 2007; Rodríguez-Herrera et al., 2014a; Reid & Gómez-Zamora, 2022; Mora & Ruedas, 2023). In this contribution, the terms *zorro pelón*, opossum, and to a lesser extent, *zarigüeya* are used to refer to *D. marsupialis*. Each of these names has a particular meaning in different contexts. *Zorro pelón*, a vernacular name specific to Costa Rica, is often used to highlight cultural and regional aspects. Opossum is a scientific term and a common name designated by Native American language and later adapted by European settlers in North America. *Zarigüeya* is the Spanish translation of opossum and is occasionally used to refer to its recognition in the Spanish-speaking world. The choice of each term in this text reflects the varied perspectives and contexts.

The importance of marsupials among the peoples of Costa Rica represents an opportunity to recover knowledge about the ecological characteristics of this group. These mammals exhibit very striking traits, which have become integral to the folkloric narrative in oral traditions. Examples of these traits include a short gestation period, rapid sexual maturity, and short lifespan, as well as adaptability to altered environments, defensive behaviors, and tolerance to venom (Wainwright, 2007; Monge-Meza & Linares-Orozco, 2009; Voss & Jansa, 2012; Reid & Gómez-Zamora, 2022). Intriguingly, in the scientific sphere, these traits have been little studied. Until 2014, scientific articles on marsupials accounted for less than 2% of the total publications for the class Mammalia in Costa Rica (Rodríguez-Herrera et al., 2014b).

Research on the ecology of marsupials in Costa Rica has been scarce for decades. Key studies conducted before 1980, such as those by Valerio (1969), Vaughan & Hawkins (1969), and Salas-Durán (1974), were followed by a long period of limited scientific activity until the works of Monge-Meza & Linares-Orozco in 2009, and subsequently by Acosta-Chaves et al. in 2018 and Muñoz-López et al. in 2022. In the field of parasitology, publications have also been sporadic, with notable exceptions such as those by Zeledón et al. (1970), Chinchilla et al. (2015), and Buhr et al. (2018). In Costa Rica, a relationship has been established between *zorro pelón* and Chagas disease because of its proximity to rural homes (Zeledón, 1981; Zeledón et al., 2005). In the field of conservation, *D. marsupialis* has received little attention, despite it being one of the mammals most frequently hit by vehicles, and it faces significant challenges in coexistence with humans (Pacheco et al. 2004; Monge-Nájera, 2018; Mora & Ruedas, 2023).

Research on the ecology of neotropical marsupials, including diet, spatial segregation, movement patterns, ecological interactions, reproductive patterns, and distribution, has maintained a constant publication rate on a regional scale (O’ Connell, 1989; Vieira & Izar, 1999; Giarla & Jansa, 2014; Barros et al., 2016; Kuhnen et al., 2017). Moreover, several studies have highlighted the resistance of marsupials to pathogens and venom. For instance, two species of didelphids and one species of *Metachirops* are resistant to different variants of the yellow fever virus (Laemmert, 1946). Vellard (1945, 1949) conducted the first study on the immunity of marsupials to the venom of viperines. After several decades, there has been a surge in the research on snake predation and immunity (Neves-Ferreira et al. 1997; Almeida-Santos et al., 2000; Gómez-Martínez et al., 2008; Voss & Jansa, 2012; Komives et al., 2016; Drabeck et al., 2020).

The only research on the ethnobiology of marsupials has focused on the alluvial plain of the Amazon in Abaetetuba, Pará, Brazil. Barros & De Aguiar (2014) found that the Gambá *D. marsupialis* has significant cultural importance to the riverine communities in Abaetetuba, as they use it as a food and medicinal resource. In this contribution, we used a methodology similar to that used by those authors to fill some gaps in knowledge about the ecology of marsupials. We present the results of a study focused on the knowledge, use, perception, and human-zorro pelón interactions of individuals from different zones (rural, semiurban, and urban) in the seven provinces of the country. Additionally, we discuss the cultural aspects of didelphids among the communities of farmers in the 19th century and indigenous peoples.

## METHODOLOGY

### Surveys and other sources of information

In the framework of two courses on mammalogy taught at the Universidad Nacional during the second semester of 2020 and the first semester of 2021, we conducted surveys on knowledge, use, perception, and human-zorro pelón interactions in three zones of Costa Rica: urban, suburban, and rural. We characterized each region using the UN-Habitat’s online page as a reference, which considers three variables: population density, type of economic activities, and vegetation cover. After validating the questionnaire, participants were requested to participate through social media. We also developed assisted surveys for older adults who did not use social networks. In this case, the student informed the participants about the use of their data and explained the questions. We used Google Forms to record participants’ responses. The questionnaire structure was semi-structured, with open and closed sections to document the knowledge and use of the *zorro pelón* and closed sections to measure the interactions and perceptions. For interactions and perceptions, we used five elements of the Likert scale: (1) very bad/harmful, (2) somewhat bad/harmful, (3) neutral, (4) somewhat good/beneficial, and (5) very good/beneficial.

As a complementary part of this study, key informants, including two professional herpetologists, a former hunter, and two elderly individuals from the Central Valley, were interviewed. Additionally, a historical document review was conducted to address the cultural aspects of *zorro pelón* and other marsupials among 19th century farmers and indigenous peoples of Costa Rica.

### Data analysis

The geographic location of respondents was georeferenced using Google Earth (2023). We then used QGIS software (2023) to create a frequency map of the surveys applied in each province. Text analysis and Likert scale element analysis were performed using R packages tidytext version 0.4.1, and likert version 1.3.5 (Bryer & Speerschneider, 2016; Silge & Robinson, 2016). Furthermore, we assessed the effect of zone type (rural, semi-urban, and urban) on the presence and absence of Likert elements in variables related to human-zorro pelón interaction and perception. This was done through one-way PerMANOVA tests using PRIMER v7 software (Clarke & Gorley, 2015). PerMANOVA tests were based on Euclidean distances, and *P* values were obtained through 999 permutations. When the type of zone was significant (*P* < 0.05) in relation to the interaction or perception variables, we conducted multiple comparisons between the levels of the predictor variables (i.e., rural vs. semi-urban, rural vs. urban, and semi-urban vs. urban). Multiple comparisons were performed using non-parametric statistical analysis, analogous to *t*-tests for multivariate analysis (pseudo-*t*). This was complemented by Monte Carlo permutations to compensate for the low number of samples (Anderson et al., 2008).

## RESULTS

### **I** Respondents’ characteristics

The surveys were conducted across 146 districts in seven provinces of Costa Rica (Fig. 1). We conducted 296 surveys with individuals from three types of zones: rural, urban, or semi-urban. Participants’ ages ranged from 18 to 75 years (Table 1). More than one-third of the respondents (39%) resided in agricultural or livestock areas, predominantly located in rural and semi-urban zones. The vast majority of respondents (99%) reported living with pets, such as cats and dogs, or raising birds and livestock. The most frequently reported occupations among the participants were students, teachers, homemakers, farmers, engineers, conservation, and tourism (Fig. 2).

**Figure 1.**
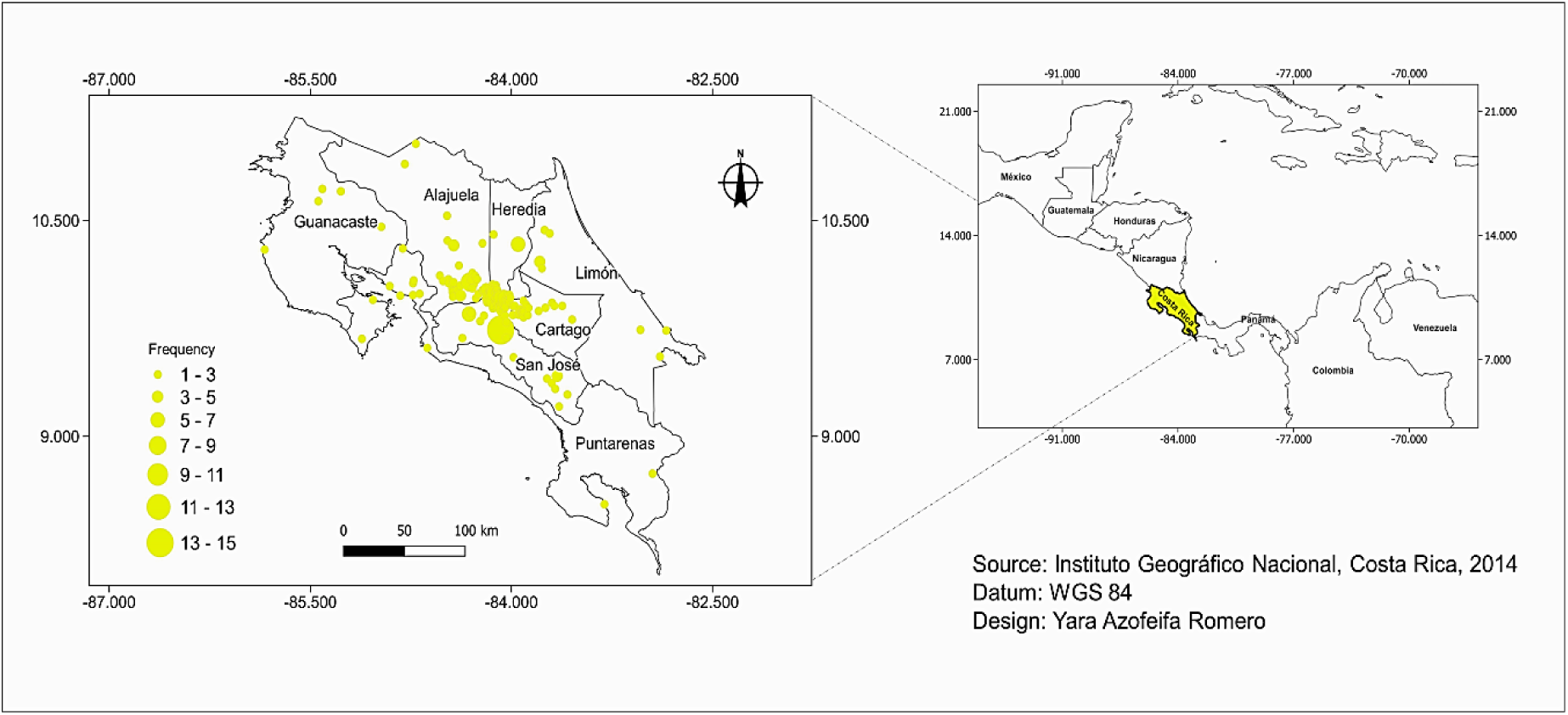
Distribution map of the 296 surveys carried out in the seven provinces of Costa Rica

**Figure 2.**
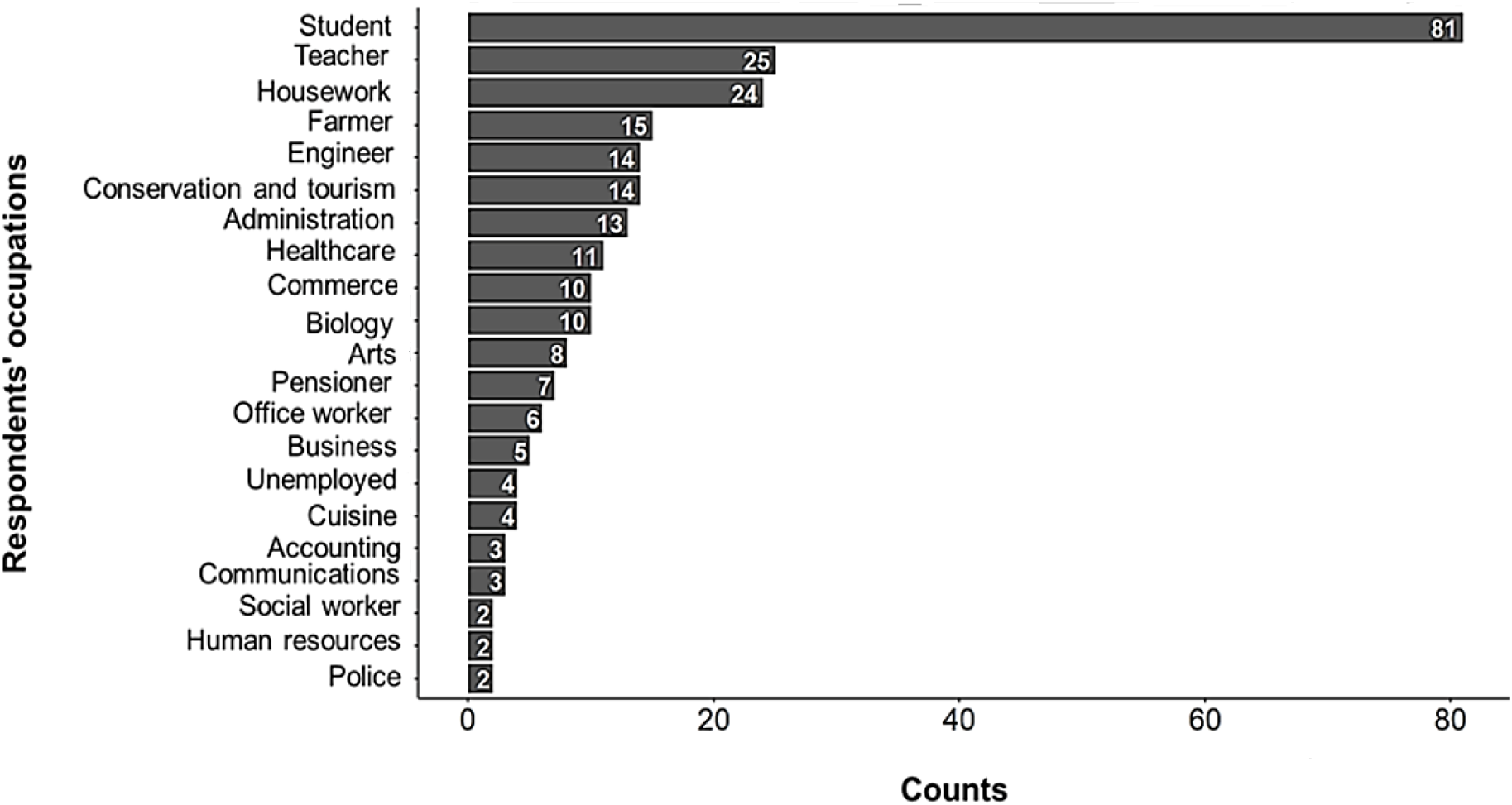
Observed frequencies of the recorded occupations for 296 people surveyed. Frequencies > 2 are included.

**Table 1.** Number of surveys carried out in each type of zone and for each age group.

| Age group | Rural | Semiurban | Urban |
| --- | --- | --- | --- |
| < 18 | 3 | 6 | 0 |
| 18-24 | 30 | 18 | 15 |
| 25-34 | 34 | 32 | 28 |
| 35-44 | 15 | 11 | 23 |
| 45-54 | 11 | 12 | 22 |
| 55-64 | 8 | 10 | 8 |
| 65-74 | 2 | 4 | 1 |
| 75+ | 1 | 1 | 1 |
| Total | 104 | 94 | 98 |

### **II** Taxonomic identification, ecology, and behavior of the zorro pelón

A majority of the individuals surveyed (83.5%) taxonomically identified *D. marsupialis* as marsupial, while others classified this mammal as rodent (15.2%) or primate (0.3%). Among the participants, 12.8% reported that they had never seen a *zorro pelón* and were, therefore, redirected to the perception section of the questionnaire. They were not asked any further questions regarding their knowledge, use, and interactions with the species. The remaining participants primarily characterized the species based on their hairless tail, snout, and fur, unattractive appearance, resemblance to a rat, behavior of “feigning death,” foul odor, and screeching noises. A few individuals mentioned the *zorricí Marmosa* sp., noting its similarity to mice. When asked to identify *D. marsupialis* from three photos showing a lesser anteater, rodent, and zorro pelón, 98% were successful.

The most frequently recorded observation sites for *D. marsupialis* were tree branches (28.5%), ground (26.7%), and house roofs (19.5%). Other reported observation sites included tree hollows, electrical wiring, and caves (Fig. 3). Regarding behavior, 66.3% of the participants indicated that this marsupial could be observed year-round. However, others noted more frequent observations or sounds during the rainy season (20.5%) or dry season (13.2%). It has been commonly reported that the nighttime is the peak activity period for this mammal (Fig. 4). Some individuals recorded that the *zorro pelon* was most active between 7 PM and 11 PM; however, two elderly individuals suggested that this initial activity period could be influenced by the presence of dogs.

**Figure 3.**
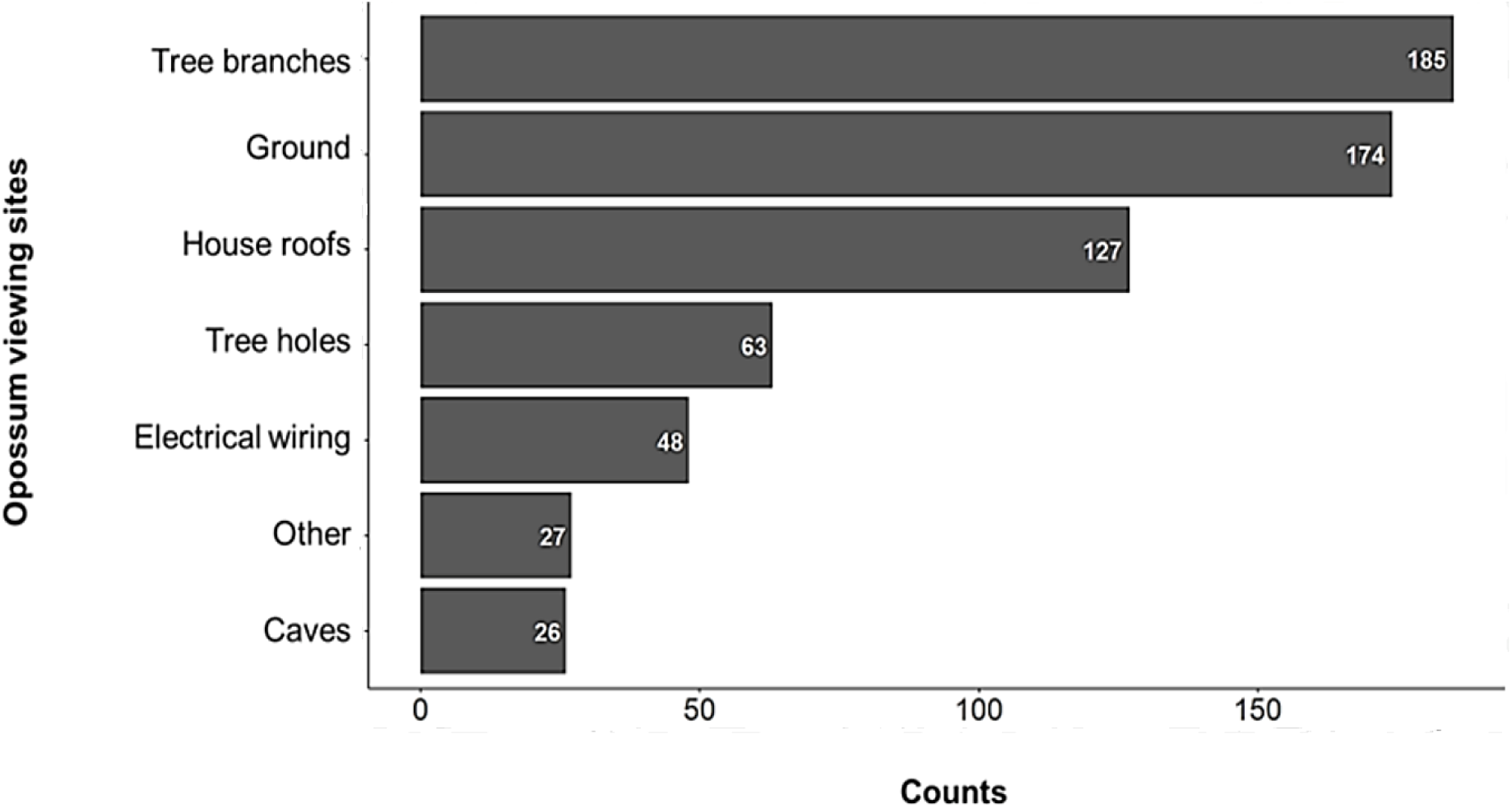
Observed frequencies of opossum viewing sites recorded by 258 people surveyed. The greater frequency with respect to the number of people surveyed was due to the possibility of marking several options in the questionnaire. Frequencies > 0 are included.

**Figure 4.**
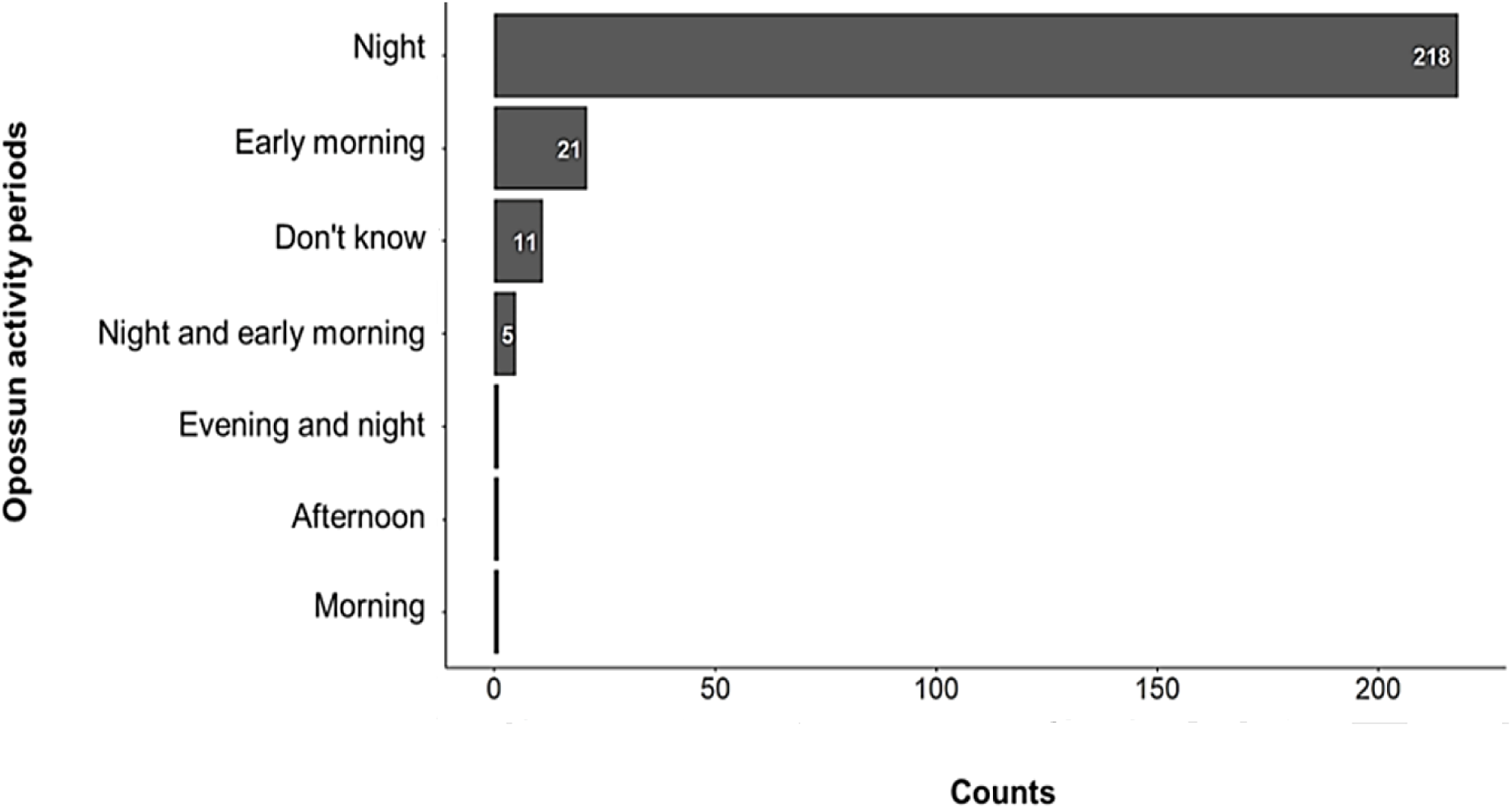
Observed frequencies of the opossum activity period recorded by 258 people surveyed. Frequencies > 0 are included.

Regarding the diet of *D. marsupialis*, the frequency of items selected by the individuals surveyed was as follows: fruits (23.9 %), animals (20.2 %), arthropods (16.8 %), food residues (15.3 %), plants (10.1 %), grains (8.0 %), carrion (4.7 %), and vegetables (0.9 %). Notable comments included: 1) fruits such as guineos, bananas, plantains, pitanga, mango, and avocado; 2) vertebrates, such as chickens, including their eggs and snakes; and 3) human and domestic food waste, such as dog and cat food. During this study, the first author observed the predation of two common snake species in the Central Valley, *Enulius flavitorques* and *Ninia maculata*, identified through photographs by two herpetologists. The notes on these predation events state: “I saw a *zorro pelón* eating something that it was manipulating. When it ran away, I approached, saw ground snakes, collected them for photography, and then released them.” Later, a herpetologist confirmed that the *zorro pelón* preys on *Ninia* sp., *Geophis* sp., and *Rhadinea* sp., as well as juvenile venomous snakes *Porthidium nasutum* and terciopelo *Bothrops asper* (J Vega Coto, Refugio de Vida Silvestre Lapa Verde, Costa Rica, personal communication). Additionally, during fieldwork, both authors observed opossums lurking around mist nets, suggesting that this mammal could be an opportunistic predator of bats. In consultation with a former *zorro pelón* hunter about other potential food items, he mentioned ceasing to hunt this marsupial for consumption after finding a toad in its stomach.

To gather opinions on population trends, we inquired about the number of *zorros pelones* observed in local areas over the past 10 years. Most of the individuals surveyed reported a reduction in the number of animals observed (Fig. 5). Possible causes of these population declines include human cruelty, dog attacks, vehicle collisions, and urban expansion.

**Figure 5.**
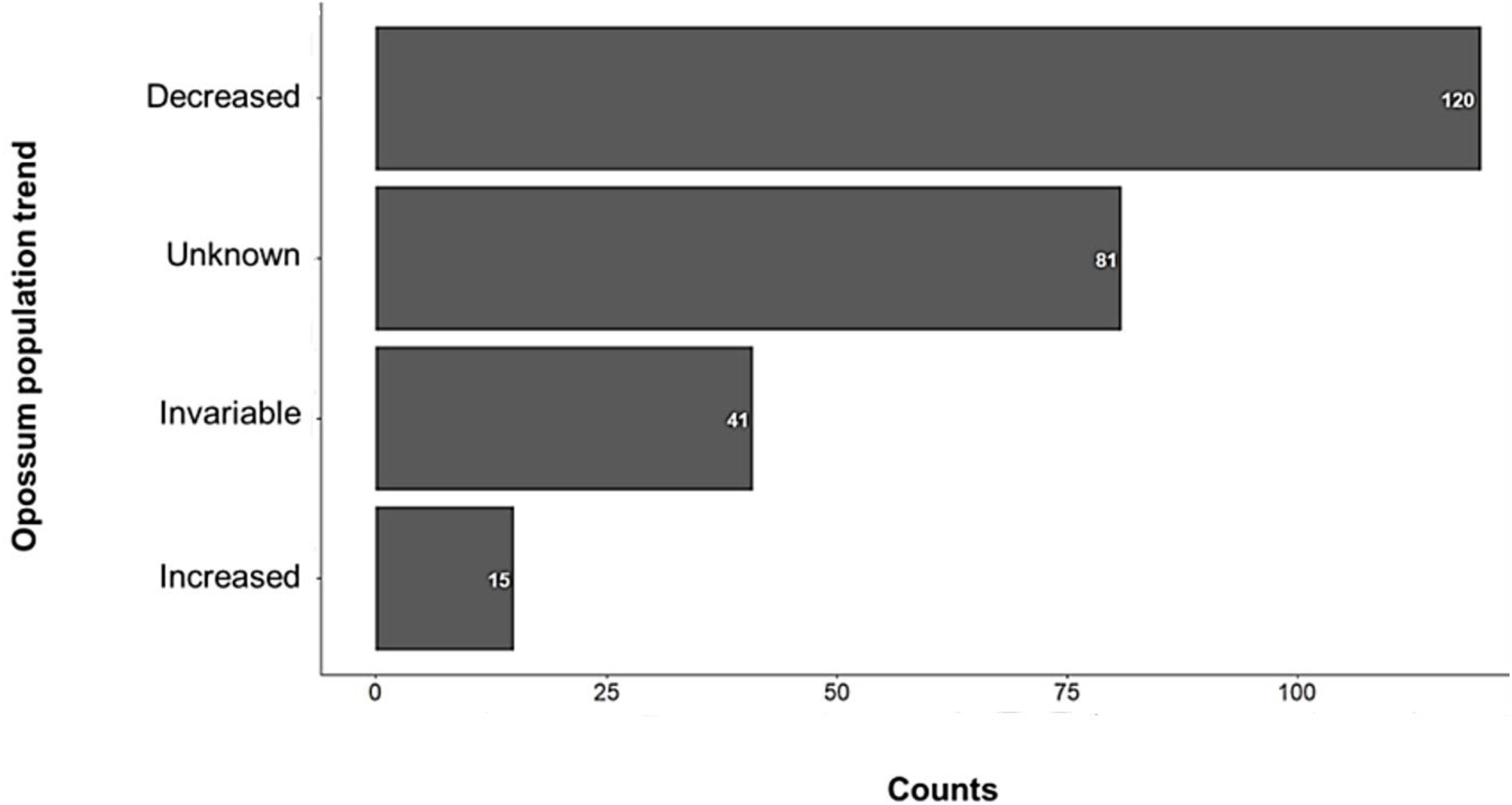
Frequencies observed in the opossum population trend recorded by the 257 people surveyed. Frequencies > 0 are included.

### **III** Interactions and perception of people towards the zorro pelón in rural, semi-urban and urban areas of Costa Rica

The likelihood of observing *D. marsupialis* varied among the individuals surveyed in the three zones (*F*_2, 257_ = 4.56, *P* = 0.001). In rural and semi-urban areas, this probability is higher than in urban areas, where 51% of city residents reported it is not very likely or not at all likely to observe this marsupial. Statistical analysis showed significant differences between the semi-urban and urban zones (*t* = 2.63, *P* = 0.001) and between the urban and rural zones (*t* = 2.45, *P* = 0.001), but not between the semi-urban and rural zones (*t* = 0.93, *P* = 0.45). While the frequency of observation differed slightly among these zones (*F*_2, 256_ = 2.49, *P* = 0.035), the only notable difference was between urban and rural areas (*t* = 2.11, *P* = 0.02). The most common frequency category across all zones is “rarely,” with a progressive increase from rural (58.7 %), to semi-urban (67.1 %), and urban zones (70.6 %). The category of observing *zorros pelones* “1-2 times per week” follows, but with a decreasing trend from rural (31.5 %) to semi-urban (20.7 %), and urban zones (12.9 %; Fig. 6).

**Figure 6.**
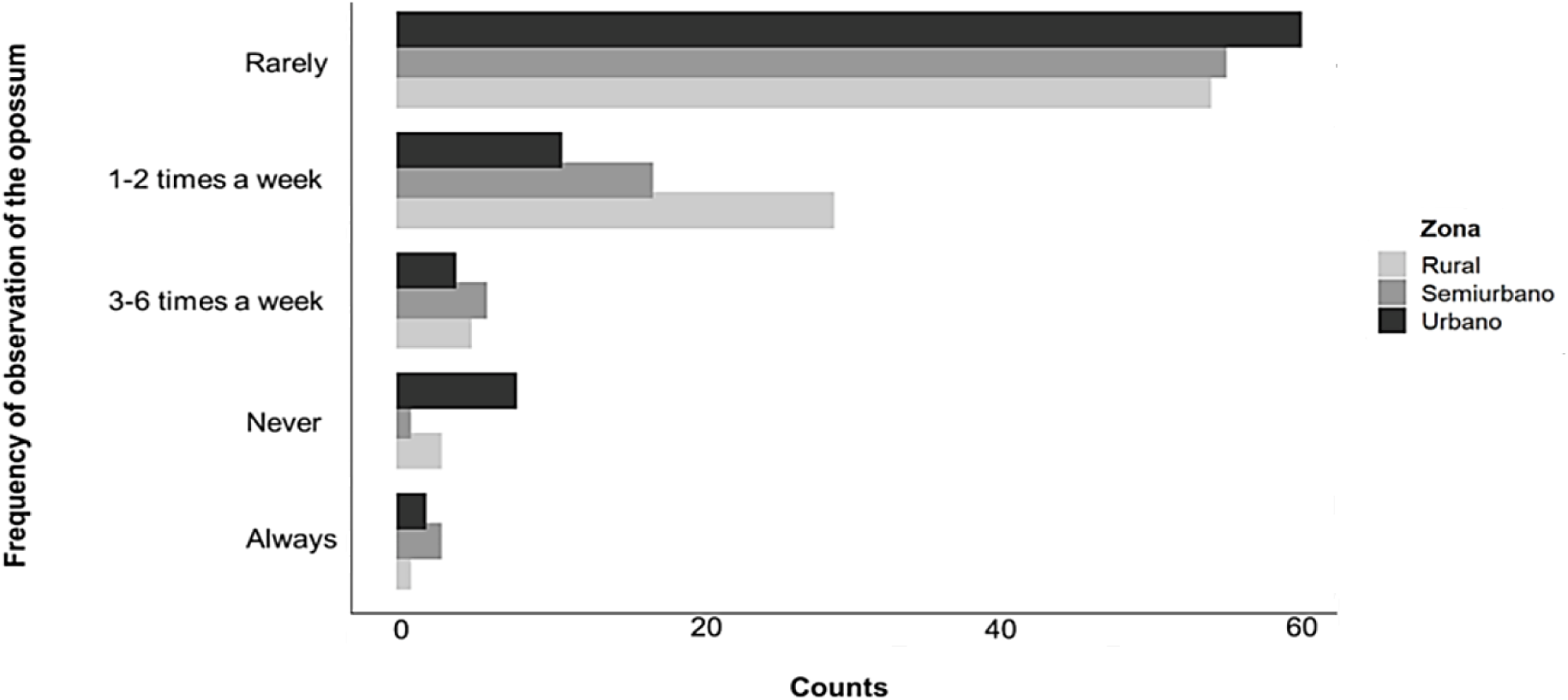
Frequencies of opossum observation recorded by 92, 82, and 95 people surveyed from rural, semi-urban and rural areas, respectively. Frequencies > 0 are included.

Regarding actions taken when encountering a *zorro pelón*, the most frequent responses, regardless of the zone, were not interrupting the animal’s activity (56.1%), scarring it away (21.6%), improving infrastructure (12.2%), relocating the animal (6.8%), killing it (2.3%), and calling the SINAC (Sistema Nacional de Áreas de Conservación) or firefighters (1.1%). Interactions with pets, particularly domestic dogs, are the most common and pose a significant danger to marsupials.

Overall, the perception of the individuals surveyed towards the *zorro pelón* tended to be more positive than negative (Fig. 7). However, in addition to its potential role as a seed disperser (*F*_2, 293_ = 2.84, *P* = 0.019), which differed significantly between semi-urban and urban zones (*t* = 1.97, *P* = 0.011), and urban and rural zones (*t* = 1.93, *P* = 0.023), other perceptions did not show significant variation between the different types of zones. It is important to note that the domestication of the *zorro pelón* was viewed negatively by the participants.

**Figure 7.**
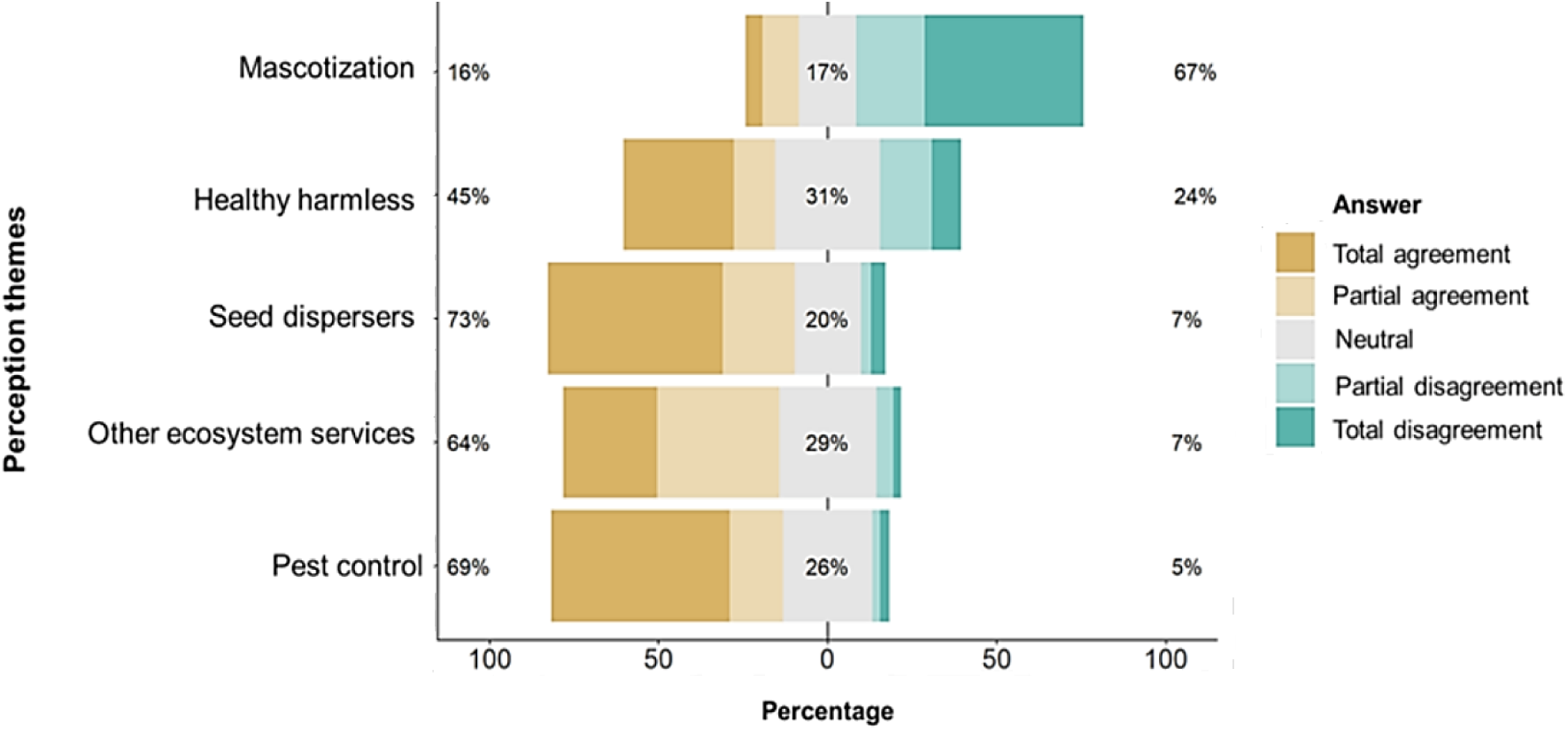
Likert scale for five questions related to perception issues of the zorro pelón from the 297 people surveyed.

### **IV** Oral tradition stories and uses of the zorro pelón in Costa Rica

The term *zorro pelón* has long been used in Costa Rica to refer to members of the order Didelphimorphia. This name likely dates back to colonial times when, due to the limited understanding of this distinctly American group, the European fox was the closest point of reference for our ancestors, despite being from a very different mammalian group. This confusion was further exacerbated by the presence of a “true” fox in Costa Rica, the gray fox *Urocyon cinereoargenteus*, which was previously mistaken for other mammals such as “tigrillo” by farmers. Interestingly, with the advent of television, the Internet, and educational channels, the term *zarigüeya* has become increasingly recognized by Costa Ricans. On one occasion, we were surprised to have to clarify that when we refer to *zorro pelón*, we are also referring to the *zarigüeya* or opossum.

Among the farmers in Costa Rica, *zorro pelón* is one of the most familiar species. Its visits to homes and its presence in people’s daily lives were so frequent that the German naturalist von Frantzius (1881) wrote about it: “The first tropical animal with whom the newly arrived foreigner establishes relationships, albeit an unpleasant one. In cities, few houses are not visited by this ugly animal. Frequently, the foreigner is awakened in the silence of the night by the extraordinary noise made by these animals with their runs and clattering steps on the thin boards of the flat roofs of the rooms, or by their visits to the pantry and kitchen, where they knock over and break plates, bowls, and other crockery; hence some foreigners, fearing that the noise is caused by burglars, get up and grab their weapons. The next day they ask for an explanation of the unexpected uproar and receive this reply: Ah, sir, it’s just the fox.”

It is believed that “fox”’ hot blood, fat, and broth can cure asthma. The meat of the *Didelphis* genus of “foxes” has been consumed and its flavor is even compared to chicken. According to participants and key informants, the consumption of *zorro pelón* meat in bars, where alcohol is consumed, was quite common in the Central Valley, and families at that time prepared it as soup and rice. These practices are likely to continue in several regions of the country. Today, as in the past, opossums are hunted because they prey on chicken (Fig. 8).

**Figure 8.**
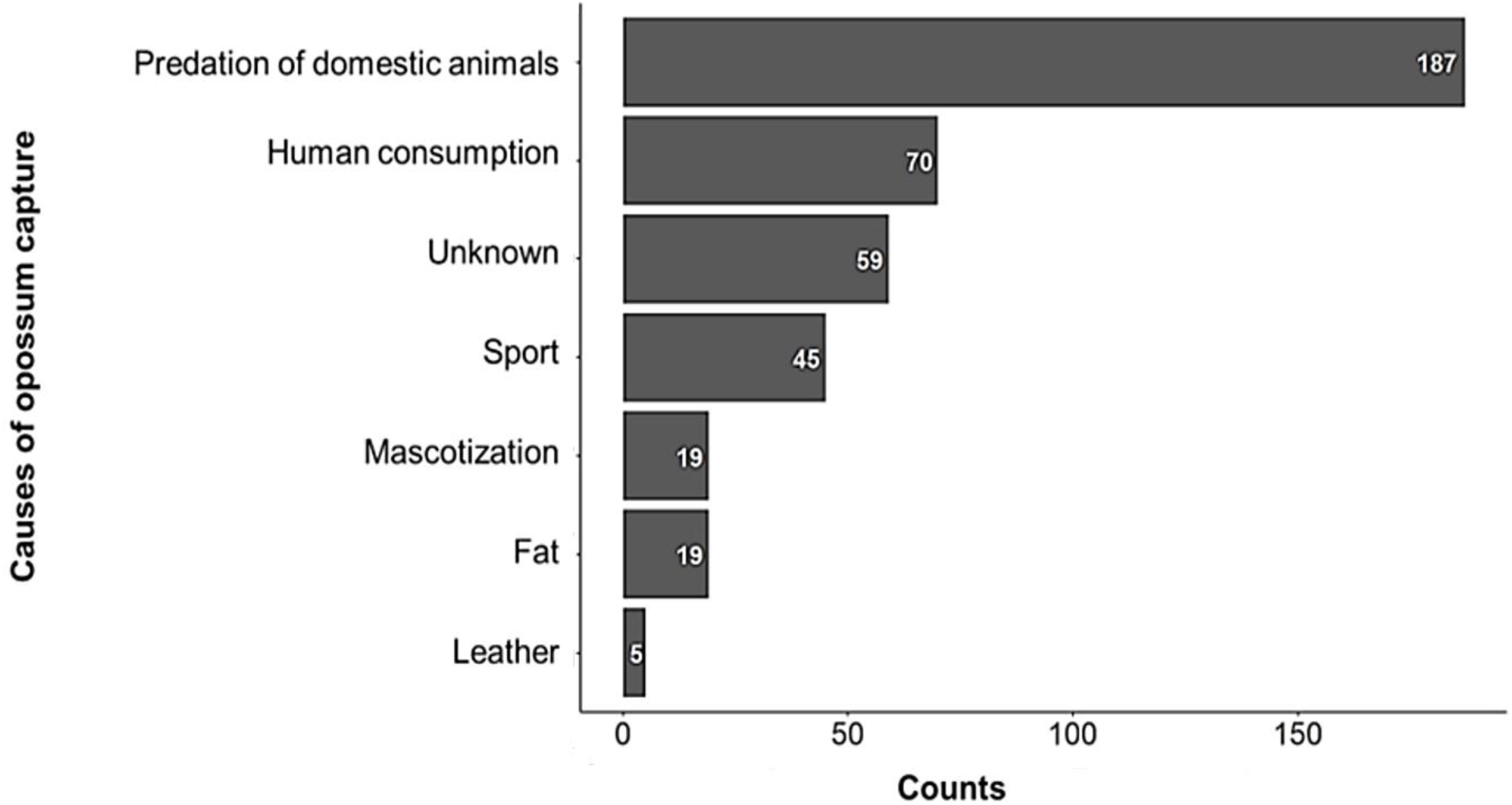
Observed frequencies of the causes of opossum capture recorded by 258 people surveyed. The greater frequency with respect to the number of people surveyed was due to the possibility of marking several options in the questionnaire. Frequencies > 0 are included.

The saying *hacerse el zorro* refers to a sleepy person who does not want to get up. This expression seems to originate from the defensive technique of the “fox” of the genus *Didelphis*, which simulates death, as mentioned earlier. Another less common saying is *más duro que un zorro pelón*, which is possibly related to the beatings this species receives, as implied by the common expression *moler un zorro a palos*. Following these beatings, the *zorro pelón* would often leave, even with several broken bones. This resilience led people to believe it was very resistant to blows (Valerio, 1969). Regarding this, one participant commented that farmers in the southern region catch them with traps, put them in bags, beat them, sometimes because they believe “foxes” eat chickens and other times for consumption.

### **V** Cultural aspects about didelphids in the indigenous peoples of Costa Rica

The *zorro pelón* was likely well known among the pre-Columbian cultures in Costa Rica, as suggested by some highly stylized pre-Columbian artifacts. These artifacts depict animals with a prominent snout and long tail, sometimes carrying one or two offspring, indicating a relationship with didelphid species. However, in cases where only one offspring is shown, this might also represent an ant bear.

Among the stories of the indigenous people of Talamanca in Costa Rica, there is a series about the rivalry between Sibö or Sibu (God) and a major devil (Sórkula, Sorblu’, or Kulerpa), which features the figure of the *zorro pelón* (*D. marsupialis*). In one of these stories, Sibu resurrects a *zorro pelón*, which then starts playing maracas (Bozzoli 2017). This action terrified the devils so much that they swore never to eat the meat of the *zorro pelón*. This story possibly led to a dietary taboo among the Talamanca people against consuming *zorro pelón*, as the animal symbolizes a transient state between life and death (Guevara, 1988), a concept likely linked to the opossum’s defensive technique of playing dead. Because of this story, the person responsible for speaking on behalf of the people to the high priest is called bikili’ (*zorro pelón*, ally of Sibu). In funerals, the opossum is represented by the singer’s maraca (Jara and García, 2003).

*Zorro pelón* also plays an important role in indigenous medicine. The Talamanca people have historically used their skin to treat illnesses related to abortion and to address improper burials (Castillo and Borge 1995). Similarly, the skins of the gray four-eyed opossum (*Philander melanurus*) and the water opossum (*Chironectes minimus*) are considered medicinal in Talamanca traditions. Carl Bovallius (1882) documented an indigenous burial in Talamanca, noting that among the items placed beside the corpse was an amulet made from *zorro pelón* skulls (Zeledón, 1997).

Within the inherited indigenous culture of the country, there are various toponyms in the names of towns and others that originate from indigenous words. Dikori is the name of a gorge in the province of Cartago. According to Quesada (2007), the word is of cabécar origin (a native tribe of Costa Rica) and means “water fox river,” an undoubted connection with *C. minimus*, which is strongly associated with aquatic environments.

### **VI** Discussion and conclusions

The opossum, a common species in Costa Rica, inhabits a diverse range of environments including rural, semi-urban and urban zones, and secondary forests, from sea level to altitudes as high as 2,850 meters (Carvajal and Aguilar, 2022, UTN-SINAC, commun. Pers). The widespread occurrence of *zorro pelón* has facilitated the collection of ecological data through public surveys, which corroborates the findings in the scientific literature. Thus, for example, the knowledge that people draw from their memories seems to approach the general aspects of *zorro pelón’s* natural history. There are observations of seasonal patterns of activity, with increases during the rainy season, which could be related to the species’ reproductive activity (O’Connell, 1989). Public knowledge about the diet of zorro pelón, including its predation on snakes and consumption of toads, often aligns with or even surpasses scientific reports (Wainwright, 2007; Acosta-Chaves et al., 2018). Regarding population trends, participants observed a decline in *zorro pelón* numbers, likely due to urban expansion. This results in habitat loss and increased contact with humans and their domestic animals, particularly dogs, which compete and prey upon or kill the opossum. Hunting and road accidents have also contributed to this decline. Nevertheless, comprehensive data to support these observations are still needed.

The image of this omnivore is associated with residing on house roofs and preying on chicken. The *zorricí Marmosa* sp. probably also occupied dwellings; however, since it is confused with mice, it would not be surprising if, in rural and semi-urban areas, people had eliminated them from their homes and caused extirpation. The great adaptability of the *zorro pelón* and *zorricí*, including the gray four-eyed opossum, *P. melanurus*, in these two zones coincides with the reports of Valerio (1969), Vaughan & Hawkins (1969), and Monge-Meza & Linares-Orozco (2009). Gathering ecological information about lesser-known species is challenging because of the general lack of taxonomic knowledge among the public.

The scientific knowledge of marsupials in Costa Rica remains limited. Natural history observations and home range studies have been conducted on the zorro de balsa or woolly opossum *Caluromys derbianus* (Salas-Durán, 1974; Muñoz-López et al. 2022). Acosta-Chaves et al. (2018) reported the feeding behavior of the *zorricí* or mouse opossum *Marmosa zeledoni*, a highly cryptic arboreal species. Little is known about the *zorricí pardo* or brown four-eyed opossum (*Metachirus myosuros*), with information only available on collection locations (McPherson, 1986). Regional texts provide valuable distribution data for these mammals in areas such as La Selva Biological Station-Braulio Carrillo, Piedras Blancas National Park, and Palo Verde National Park (Landmann et al., 2008; Stoner et al., 2004; Timm, 1994; Timm et al., 1989). Timm’s text and other authors from 1989 are particularly detailed, listing species with notes on aspects such as distribution, specimens examined from collections, and sightings in the area. However, more work is needed to understand the ecology of this group.

The use of opossums in food and medicinal practices in various communities, including modern non-indigenous populations and indigenous groups in Costa Rica and Brazil, is intriguing. This practice, involving the utilization of opossum meat, blood, and fat, has been maintained throughout various cultures and continues in contemporary times (Barros & De Aguiar, 2014). In the Amazon region of Brazil, Gambá *D. marsupialis* has been documented in traditional medicine to treat conditions such as arthritis, asthma, sore throat, and inflammation (Barros & Azevedo, 2014). This medicinal application is quite similar to what we report in our study, so it would not be surprising if these practices are repeated in other neotropical regions. Although the opossum acts as a reservoir of pathogens, and its consumption could have implications for human health (Villagra-Blanco et al., 2019), until now, the only thing that has been addressed is the health of opossums in a state of captivity (VandeBerg et al., 1986; Cothran et al. 1990).

The results of these surveys require careful analysis, particularly concerning perceptions and interactions, since the significance of the responses could vary with increased survey effort and greater inclusion of rural participants. While the general public’s perception may not be overly negative, there has been a notable devaluation of marsupials over time compared to indigenous cultures’ views. Physical attacks on these mammals, for example, are not linked to indigenous practices and may stem from colonial influence. This should be considered in conservation strategies, particularly in areas that have reported population reductions. Efforts should be made to promote appreciation and dispel negative stereotypes, for example, by informing people of their ecological role and their ability to resist pathogens and venoms. In this context, the rich symbolism of the opossum can be leveraged to adapt the message to the various regions of America.

## AGRADECIMIENTOS

To the students of the mammal course who contributed to the design, validation, and implementation of the project: Pablo Gutiérrez Campos, Natalia Alfaro Mejía, Alexander Monge Jiménez, Verónica Orozco Fernández, Rebeca Solano Gómez, Oscar Brenes Hernández, Yaudi Castillo Pérez, Raquel Castro Lara, Yalendi Corrales Vargas, Adrián Rodríguez Salazar, Marilyn Ureña Salazar, Karen Valverde Arias, Aaron Vargas Briseño, Sharys Benavides Guido, Ana Abarca Méndez, Daniela Segura Astúa, Genesis Rodríguez Naranjo, Jasmín Mejía Vargas, Julio Zúñiga Marín y Sara Lizano Conejo. We express our deepest gratitude to Francisco Javier Flórez Oliveros for providing us with the opportunity to contribute to this compendium. We harbor the firm hope that his dedicated efforts towards the conservation of marsupials will unite the people of America.

## Supporting information

Supplementary Material_Survey_The opossum and other didelphids

## REFERENCES

1. Acosta-Chaves, V. J., Sosa-Bartuano, Á., Morera-Chacón, B. H., & Jiménez-Castro, J. E. (2018). Records of preys hunted by the Zeledon’s Mouse Opossum Marmosa zeledoni Goldman, 1911 (Didelphimorphia: Didelphidae) in Costa Rica. Food Webs, 16, e00094. 10.1016/j.fooweb.2018.e00094

2. Almeida-Santos, S. M., Antoniazzi, M. M., Sant’Anna, O. A., & Jared, C. (2000). Predation by the opossum *Didelphis marsupialis* on the rattlesnake *Crotalus durissus*. Current Herpetology, 19(1), 1–9.

3. Anderson, M. J., Gorley, R. N., & Clarke, R. (2008). PERMANOVA+ for PRIMER: Guide to software and statistical methods. PRIMER-E.

4. Barros, F. B., & De Aguiar Azevedo, P. (2014). Common opossum (Didelphis marsupialis Linnaeus, 1758): Food and medicine for people in the Amazon. Journal of Ethnobiology and Ethnomedicine, 10(1), 65. 10.1186/1746-4269-10-65

5. Barros, C. S., Martins, T. K., Puettker, T., & Pardini, R. (2016). Long distance and short time movement of a small neotropical marsupial. Oecologia Australis, 20(3), 75–79. 10.4257/OECO.2016.2003.09

6. Bozzoli, M. E. (2017). En torno al tema del gallo resucitado en relatos bribris, Costa Rica. Cuadernos de Antropología, 27(1), 1–18. 10.15517/cat.v27i1.29061

7. Bryer, J., & Speerschneider, K. (2016). Likert: Analysis and Visualization Likert Items. R package version 3.6.3.

8. Castillo, V. R., & Borge, C. (1995). Especies de flora y fauna usadas por los indígenas Bribrís y Cabecares de Talamanca. Proyecto Ecología Cultural de Talamanca y Comisión para la Defensa de los Derechos Indígenas de Talamanca.

9. Chinchilla, M., Valerio, I., & Duszynski, D. (2015). Endogenous life cycle of *Eimeria marmosopos* (Apicomplexa: Eimeriidae) from the Opossum, *Didelphis marsupialis* (Didelphimorphia: Didelphidae) in Costa Rica. Journal of Parasitology, 101(4), 436–443. 10.1645/15-730.1

10. Clarke, K.R., & Gorley, R.N. (2015). PRIMER v7: User Manual/Tutorial [Manual]. PRIMER-E.

11. Cothran, E. G., Haines, C. K., & VandeBerg, J. L. (1990). Age effects on hematologic and serum chemical values in gray short-tailed opossums (Monodelphis domestica). Laboratory Animal Science, 40(2), 192–197.

12. de Buhr, N., Bonilla, M. C., Jiménez-Soto, M., von Köckritz-Blickwede, M., & Dolz, G. (2018). Extracellular trap formation in response to Trypanosoma cruzi infection in granulocytes isolated from dogs and common opossums, natural reservoir hosts. Frontiers in Microbiology, 9, 966. 10.3389/fmicb.2018.00966

13. Drabeck, D. H., Rucavado, A., Hingst-Zaher, E., Cruz, Y. P., Dean, A. M., & Jansa, S. A. (2020). Resistance of South American opossums to vWF-binding venom C-type lectins. Toxicon, 178, 92–99. 10.1016/j.toxicon.2020.02.024

14. Jara-Murillo, C. V., & García, A. (2003). Diccionario de Mitología Bribri. San José, C.R.: Editorial Universidad de Costa Rica (EUCR).

15. Google LLC. (2023). Google Earth (Versión 7.3.6) [Software]. Disponible en https://www.google.com/earth/

16. Gómez-Martínez, M. J., Gutiérrez, A., & DeClerck, F. (2008). Four-eyed opossum (*Philander opossum*) predation on a coral snake (*Micrurus nigrocinctus*). Mammalia, 72(4), 350–351. 10.1515/MAMM.2008.031

17. Giarla, T., & Jansa, S. (2014). The role of physical geography and habitat type in shaping the biogeographical history of a recent radiation of Neotropical marsupials (Thylamys: Didelphidae). Journal of Biogeography, 41(8), 1547–1558. 10.1111/jbi.12320.

18. Komives, C. F., Sanchez, E. E., Rathore, A. S., White, B., Suntravat, M., Balderrama, M., Cifelli, A., & Joshi, V. (2017). Opossum peptide that can neutralize rattlesnake venom is expressed in *Escherichia coli*. Biotechnology Progress, 33(1), 81–86. 10.1002/btpr.2386

19. Kuhnen, V., Romero, G., Linhares, A., Vizentin-Bugoni, J., Porto, E., & Setz, E. (2017). Diet overlap and spatial segregation between two neotropical marsupials revealed by multiple analytical approaches. PLoS ONE, 12. 10.1371/journal.pone.0181188.

20. Laemmert, H. W. J. (1946). Studies on susceptibility of Marsupialia to different strains of yellow fever virus. The American Journal of Tropical Medicine, 26(1), 33–46. doi:10.4269/ajtmh.1946.s1-26.33

21. Landmann A, Walder C, Vorauer A, Emser T. 2008. Mammals of the Piedras Blancas National Park, Costa Rica: species composition, habitat associations and efficiency of research methods – a preliminary overview. Stapfia 88: 409–422.

22. McPherson, A. B. (1986). Comments on the status of Metachirus nudicaudatus dentatus (Goldman, 1912) in Costa Rica. Brenesia, 24, 375–377.

23. Monge-Meza, J., & Linares-Orozco, J. (2009). Presencia del zorro de cuatro ojos (Philander opossum) en el cultivo de piña (Ananas comusus). Agronomía Mesoamericana, 21(2), 343–347. 10.15517/am.v21i2.4898

24. Monge-Nájera, J. (2018). What can we learn about wildlife killed by vehicles from a citizen science project? A comparison of scientific and amateur tropical roadkill records. UNED Research Journal, 10, 57–60. 10.22458/urj.v10i1.2041

25. Mora, J. M., & Ruedas, L. A. (2023). Updated list of the mammals of Costa Rica, with notes on recent taxonomic changes. Zootaxa, 5357(4), 451–501. 10.11646/zootaxa.5357.4.1

26. Muñoz-López, M., Villalobos-Chaves, D., Rojas-Valerio, E., & Rodríguez-Herrera, B. (2022). Home range and movement ecology of the woolly opossum (Caluromys derbianus) in a Neotropical rainforest of Costa Rica. Therya Notes, 3, 119–124. 10.12933/therya_notes-22-82

27. Neves-Ferreira, A. G. C., Perales, J., Ovadia, M., Moussatché, H., & Domont, G. B. (1997). Inhibitory properties of the antibothropic complex from the South American opossum (Didelphis marsupialis) serum. Toxicon, 35(6), 849–863. 10.1016/S0041-0101(96)00195-X

28. O’connell, M. A. (1989). Population Dynamics of Neotropical Small Mammals in Seasonal Habitats. Journal of Mammalogy, 70(3), 532–548. 10.2307/1381425

29. ONU-Habitat. (n.d.). ¿Cómo definir ciudades, pueblos y áreas rurales? Recuperado el 24 de diciembre de 2023, de https://www.onuhabitat.org.mx.

30. Pacheco, J., Ceballos, G., Daily, G. C., Ehrlich, P. R., Suzán, G., Rodríguez-Herrera, B., & Marcé, E. (2006). Diversidad, historia natural y conservación de los mamíferos de San Vito de Coto Brus, Costa Rica. Revista de Biología Tropical, 54(1), 219–240. 10.15517/rbt.v54i1.13998

31. QGIS Development Team. (2023). QGIS Geographic Information System. Open-Source Geospatial Foundation Project (Versión 3.32) [Software]. Disponible en: https://www.qgis.org

32. Quesada Pacheco, M. (2007). Toponimia indígena de Costa Rica. Revista de Filología y Lingüística de la Universidad de Costa Rica, 32(2), 203–259. 10.15517/rfl.v32i2.4297

33. Reid, F., & Gómez-Zamora, G. (2022). Pocket guide to the mammals of Costa Rica. Ithaca, New York: Comstock Publishing Associates, an imprint of Cornell University Press.

34. Ramírez-Fernández, J. D., Sánchez, R., May-Collado, L. J., González-Maya, J. F., & Rodríguez-Herrera, B. (2023). Revised checklist, conservation status, and endemism of the mammals of Costa Rica. Therya, 14(1), 233–244. 10.12933/therya-23-2142

35. Rodríguez-Herrera, B., Ramírez-Fernández, J. D., Villalobos-Chaves, D., & Sánchez, R. (2014a). Actualización de la lista de especies de mamíferos vivientes de Costa Rica. Mastozoología Neotropical, 21(2), 275–289.

36. Rodríguez-Herrera, B., Sánchez, R., & Alpízar, P. (2014b). Historia de la mastozoología en Costa Rica. En J. Ortega, J., L. Martínez, & D. G. Tirira (Eds.), Historia de la Mastozoología en Latinoamérica, las Guayanas y el Caribe (pp. 175–188). Editorial Murciélago Blanco y Asociación Ecuatoriana de Mastozoología.

37. Salas-Durán, S. (1974). Algunas observaciones sobre el hábito de vida del “Zorro de Balsa” Caluromys derbianus (Marsupialia, Didelphidae) en la vertiente del Pacífico de Costa Rica. ÓBios, 2, 11–15.

38. Silge, J., & Robinson, D. (2016). tidytext: Text Mining and Analysis Using Tidy Data Principles in R. JOSS, 1(3). 10.21105/joss.00037

39. Stoner, K. E., & Timm, R. M. (2004). Tropical dry-forest mammals of Palo Verde: Ecology and conservation in a changing landscape. En G. W. Frankie, A. Mata, & S. B. Vinson (Eds.), Biodiversity conservation in Costa Rica: Learning the lessons in a seasonal dry forest (pp. 48-66). University of California Press.

40. Timm, R. M., Wilson, D. E., Clauson, B. L., LaVal, R. K., & Vaughan, C. S. (1989). Mammals of the La Selva–Braulio Carrillo Complex, Costa Rica. North American Fauna, 75, 1–162.

41. Timm, R. M. (1994). Mammals. En L. A. McDade, K. S. Bawa, H. A. Hespenheide, & G. S. Hartshorn (Eds.), La Selva: ecology and natural history of a neotropical rain forest (pp. 394-397). Chicago: University of Chicago Press.

42. Valerio, C. E. (1969). La gran capacidad adaptativa del zorro pelón. Revista de la Universidad de Costa Rica, 26, 43–44.

43. VandeBerg, J., Cothran, E., & Kelly, C. (1986). Dietary effects on hematologic and serum chemical values in gray short-tailed opossums (Monodelphis domestica). Laboratory Animal Science, 36 (1), 32–6.

44. Vaughan, C., & Hawkins, L. (1969). Late dry season habitat use of common opossum, Didelphis marsupialis (Marsupialia: Didelphidae) in neotropical lower montane agricultural areas. Revista de Biología Tropical, 47, 263–269. 10.15517/RBT.V47I1-2.19075.

45. Vellard, J. (1945). Resistencia de los “Didelphis” (zarigueya) a los venenos ofídicos (Nota prévia). Revista Brasileira de Biologia, 5, 463–467.

46. Vellard, J. (1949). Investigaciones sobre inmunidad natural contra los venenos de serpientes. Publicaciones del Museo de Historia Natural de Lima, Serie A Zoología, 2, 1–80.

47. Vieira, E., & Izar, P. (1999). Interactions between aroids and arboreal mammals in the Brazilian Atlantic rainforest. Plant Ecology, 145, 75–82. 10.1023/A:1009859810148

48. Villagra-Blanco, R., Barrantes-Granados, O., Montero-Caballero, D., Romero-Zúñiga, J., & Dolz, G. (2019). Seroprevalence of *Toxoplasma gondii* and *Neospora caninum* infections and associated factors in sheep from Costa Rica. Parasite Epidemiology and Control, 4. 10.1016/j.parepi.2019.e00085

49. von Frantzius, A. (1881). Los mamíferos de Costa-Rica. En L. Fernández (Ed.), Documentos para la historia de Costa Rica, tomo 1. San José: Imprenta Nacional

50. Voss, R. S., & Jansa, S. A. (2012). Snake-venom resistance as a mammalian trophic adaptation: lessons from didelphid marsupials. Biological Reviews, 87(4), 822–837. 10.1111/j.1469-185X.2012.00222.x

51. Wainwright, M. (2007). The mammals of Costa Rica: A Natural history and field guide. Ithaca, New York, y Londres, Reino Unido: Cornell University Press.

52. Zeledón, R., Solano, G., Sáenz, G., & Swartzwelder, J. C. (1970). Wild reservoirs of *Trypanosoma cruzi* with special mention of the opossum, *Didelphis marsupialis*, and its role in the epidemiology of Chagas’ disease in an endemic area of Costa Rica. Journal of Parasitology, 56(1), 38.

53. Zeledón, R. (1981). El *Triatoma dimidiata* (Latreille, 1811) y su relación con la enfermedad de Chagas. EUNED, Costa Rica.

54. Zeledón, R., Calvo, N., Montenegro, V. M., Lorosa, E. S., & Arévalo, C. (2005). A survey on *Triatoma dimidiata* in an urban area of the province of Heredia, Costa Rica. Memórias do Instituto Oswaldo Cruz, 100(6), 507–512. 10.1590/s0074-02762005000600002

55. Zeledón, C. E. (1997). Viajes por la república de Costa Rica I. Editorial de la Dirección de Publicaciones, Ministerio de Cultura, Juventud y Deporte, Museo Nacional de Costa Rica, San José.

