## Supplementary Material_Survey_The opossum and other didelphids for "The opossum and other didelphids in the cultures of Costa Rican society and indigenous peoples: an approach to ecology and conservation"

### Survey on knowledge, perceptions, and human-opossum interactions in Costa Rica

The following survey aims to evaluate knowledge, perception, and human-opossum interactions in different regions of Costa Rica. This research is part of a collaborative project being conducted in the mammalogy course at the National University, from which a scientific note will be generated. If you agree to participate, we assume that we have your consent to use the information for a scientific purpose.

***Required**

**I. Information of the person surveyed**

If you wish to actively participate in this project, please include your name, an email address, or a phone number; if not, press continue to proceed with the interview.

________________________________________________________

2. Age *

______ < 18

______ 18-24

______ 25-34

______ 35-44

______ 45-54

______ 55-64

______ 65-74

______ 75+

3. Gender*

______ Woman

______ Man

______ Other

4. Occupation*

____________________________________________________________

5. Province, canton, and district where you have lived most of your life*

____________________________________________________________

6. Type of area where you currently live*:

_____ Rural

_____ Semi-urban

_____ Urban

7. Currently, do you raise or live with the following domestic animals (you may select multiple options)*:

_____ Poultry (chickens, roosters, chicks, geese, turkeys, ducks, etc.)

_____ Livestock (cattle, goats, sheep, horses, pigs)

_____ Rabbits

_____ Cats

_____ Dogs

_____ Other

8. If you answered 'other' in the previous question, please specify the animal; if you selected any other option, press continue to proceed with the interview.

______________________________________________________________

9. Are you currently living in an agricultural area? *

_____ Yes (skip to question 10)

_____ No (skip to question 11)

10. What crops are grown near where you live?*

______________________________________________________________________

**II. General knowledge of the respondent**

11. The opossum is:

_____ A rodent (skip to question 13)

_____ A marsupial (skip to question 13)

_____ A primate (skip to question 13)

_____ Other (skip to question 12)

12. What type of animal is the opossum?*

______________________________________________________________________

13. Have you had the opportunity to observe opossums?*

_____ Yes (skip to question 14)

_____ No (skip to question 40)

14. How do you distinguish this animal from others? If you know, include general characteristics about fur color, size, smell, and sound emitted.

15. In which image do you see an opossum?

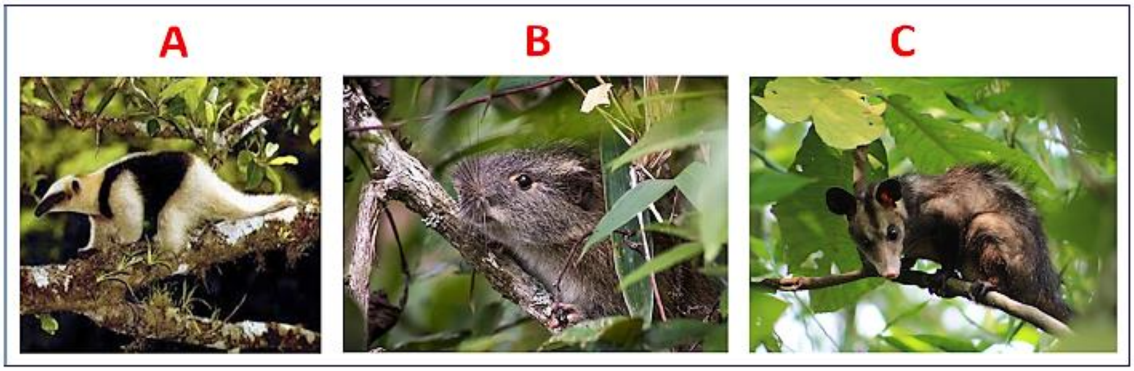


16. At what time of day do you think the opossum is most active?*

______ Early morning

______ Afternoon

______ Night

______ Night and early morning

______ Evening and night

______ I don't know

17. At what time of year do you think it is more common to see opossums?*

______ Dry season

_____ Rainy season

_____ Either season

18. Thinking about the last 10 years, do you think the number of opossums in your area*

_____ Has increased

_____ Has remained the same

_____ Has decreased

_____ Do not know

19. If you wish, you can comment on your previous answer; if not, press continue to proceed with the interview.

__________________________________________________________________

20. According to your knowledge, what is the diet of the opossum? You may select multiple options*

_____ Carnivorous (e.g., chickens)

_____ Insectivorous and related groups (e.g., insects and ticks)

_____ Herbivorous (e.g., plants)

_____ Frugivorous (e.g., fruits)

_____ Granivorous (e.g., grains)

_____ Leftovers

_____ Carrion

21. If you wish, you can comment on your previous answer; if not, press continue to proceed with the interview

_______________________________________________________________

22. In which places have you been able to observe opossums? You may select multiple options*

_____ On the ground

_____ On tree branches

_____ In tree hollows

_____ In caves

_____ On electrical wiring

_____ On house roofs

_____ Othe

23. If you wish, you can comment on your previous answer; if not, press continue to proceed with the interview

_______________________________________________________________

24. How many times a year do you think opossums reproduce?*

_____ Once a year

_____ Twice a year

_____ Three times a year

_____ More than three times a year

25. How many offspring do you think an opossum can have?*

_____ 1 - 3

_____ 4 - 6

_____ 7 - 10

_____ 11 - 15

_____ 16 – 20

26. Have you heard any oral tradition stories about opossums?*

_____ Yes

_____ No (skip to question 28)

27. What oral tradition story do you know?

________________________________________________

**III. Uses of the opossum**

28. Why are opossums captured or hunted? You may select multiple options*

_____ For family food

_____ Due to damage or predation of domestic animals

_____ Their hide is used to make clothing or other valuable items

_____ Their fat is used to make ointments/creams/salves

_____ For pets

_____ For sport

_____ I don't know the reasons

29. If you wish, you can comment on your previous answer; if not, press continue to proceed with the interview

_______________________________________________________________

30. Do you know any methods of capturing or hunting opossums?*

_____ Yes

_____ No (skip to question 32)

31. What types of methods for capturing or hunting opossums do you know?

_______________________________________________________________

32. Do you know anyone who engages in the following activities (you may select multiple options)*

_____ Breeding of the species to maintain the population

_____ Breeding for sale of offspring

_____ Breeding for meat consumption

_____ Breeding for sale of meat

_____ Training for a specific purpose

_____ None of the above

33. If you wish, you can comment on your previous answer; if not, press continue to proceed with the interview

_____________________________________________________________

**IV. Human-Opossum Interactions**

34. How likely is it for you to observe an opossum in the place where you currently live? *

_____ Very likely (skip to question 35)

_____ Likely (skip to question 35)

_____ Somewhat likely (skip to question 35)

_____ Unlikely (skip to question 35)

_____ Not at all likely (skip to question 40)

35. How often do you usually observe these animals in your place of residence?*

_____ Every day

_____ 3-6 times per week

_____ 1-2 times per week

_____ Rarely

36. What do you do when an opossum approaches the surroundings of your house?*

_____ Do not interrupt the activity of the opossum

_____ Relocate the animal on your own

_____ Scare it away

_____ Kill it

_____ Call the firefighters

_____ Call MINAE (MInisterio de Ambiente y Energía)

_____ Improve infrastructure to prevent the opossum from entering

37. Have you ever seen domestic animals (dogs, cats) interacting with opossums?*

_____ Yes

_____ No (skip to question 39)

38. If you wish, you can comment on your previous answer; if not, press continue to proceed with the interview

___________________________________________________________________

39. What was the interaction they had? Please comment*

___________________________________________________________________

**V. Opossum perception**

40. Opossums are*

_____ Extremely beneficial

_____ Beneficial

_____ Neither beneficial nor harmful (= neutral)

_____ Harmful

_____ Extremely harmful

41. If you wish, you can comment on your previous answer; if not, press continue to proceed with the interview

_______________________________________________________________

42. Do opossums act as natural controllers of other animals?*

____ Strongly agree

____ Somewhat agree

_____ Neutral

_____ Somewhat disagree

_____ Strongly disagree

43. Do opossums disperse seeds?

____ Strongly agree

____ Somewhat agree

_____ Neutral

_____ Somewhat disagree

_____ Strongly disagree

44. If you wish, you can comment on your previous answer; if not, press continue to proceed with the interview

_______________________________________________________________

45. Do you believe that opossums transmit viruses that cause diseases in humans?*

____ Strongly agree

____ Somewhat agree

_____ Neutral

_____ Somewhat disagree

_____ Strongly disagree

46. Do you know of any cases where someone has become ill after handling an opossum? If you do not know, press continue to proceed with the interview.

______________________________________________________________

47. Do you believe that opossums are beneficial for the environment?*

_____ Yes

_____ No (skip to question 49)

48. If you wish, you can comment on your previous answer; if not, press continue to proceed with the interview

______________________________________________________________

49. Provide your personal opinion on the hunting of these animals*

______________________________________________________________

50. Do you believe that opossums could be domesticated in the same way as a dog or cat?*

____ Strongly agree

____ Somewhat agree

_____ Neutral

_____ Somewhat disagree

_____ Strongly disagree

**END OF SURVEY**
